# Role of Disulfide Bonds in Membrane Partitioning of a Viral Peptide

**DOI:** 10.1101/2021.09.21.461184

**Authors:** Samapan Sikdar, Manidipa Banerjee, Satyavani Vemparala

**Affiliations:** The Institute of Mathematical Sciences, C.I.T. Campus, Taramani, Chennai 600113, India; Homi Bhabha National Institute, Training School Complex, Anushakti Nagar, Mumbai, 400094, India; Kusuma School of Biological Sciences, Indian Institute of Technology-Delhi, Hauz Khas, New Delhi - 110016, India

**Keywords:** Disulfide bond, Membrane Active Peptide, Viroporin, Membrane Remodelling, Lipid Packing Defects, Molecular Dynamics Simulations

## Abstract

The importance of disulfide bond in mediating viral peptide entry into host cells is well known. In the present work, we elucidate the role of disulfide (SS) bond in partitioning mechanism of membrane active Hepatitis A Virus-2B (HAV-2B) peptide, which harbours three cysteine residues promoting formation of multiple SS-bonded states. The inclusion of SS-bond not only results in a compact conformation but also induces distorted α-helical hairpin geometry in comparison to SS-free state, resulting in reduced hydrophobic exposure. Owing to this, the partitioning of HAV-2B peptide is completely or partly abolished. In a way, the disulfide bond regulates the partitioning of HAV-2B peptide, such that the membrane remodelling effects of this viral peptide are significantly reduced. The current findings may have potential implications in drug designing, targeting the HAV-2B protein by promoting disulfide bond formation within its membrane active region.

## 1. Introduction

Disulfide (SS) bond formation involves oxidation of thiol (SH) groups of two spatially proximal cysteine residues leading to a covalent linkage between their side-chains. The thiol-disulfide exchange reaction, regulated by oxidoreductases, like thioredoxin and protein disulfide isomerise (PDI),[1] predominantly occurs in endoplasmic reticulum and occasionally at other cellular sites.[2] The presence of disulfide bonds contribute to both protein structure and function by inducing conformational stability, facilitating protein folding and assembly, sensing changes in redox environment to modulate protein activity and localization.[1, 2] This plethora of functions illustrates the importance of disulfide bonds in both secretory and membrane proteins, like, Prion[3, 4], Src family kinases[5], Voltage-Dependent Anion Channel[6], GPCRs[7] to name a few.

Membrane active peptides often harbour multiple cysteine residues, which may remain in reduced thiol state or in oxidized disulfide linked state.[8] Several studies have highlighted the role of redox status of cysteine residues in modulating permeability of these membrane active peptides.[1, 9] For instance, antimicrobial peptides belonging to defensin family, characterized by presence of three intra-molecular SS-bridges, have been vastly studied in context of host defence mechanism against virulent microbes, by varying the number of disulfide bond and their native connectivity.[10–14] Experiments reveal enhanced antimicrobial activity upon reduction of SS-bonds of Human *β* – defensin 1 (hBD-1)[10] and hBD-4[11]. Further, molecular dynamics (MD) simulation studies of hBD-3 analogs lacking SS-linkages induce significant disruption of negatively charged lipid bilayers, compared to their native counter-part.[13] Similarly, the antimicrobial activity of other disulfide rich peptides has been known to be regulated by presence / absence of SS-bonds.[15–18]

Environment dependent thiol-disulfide switching plays important role in mediating virus entry into cells.[9] In this regard, it is required for Human immunodeficiency virus-1 (HIV-1) envelope protein dissociate into two subunits namely, gp120 and gp41, upon interaction with host cell receptors followed by reduction of redox active SS-bonds to allosterically unmask membrane active fusion peptide initiating membrane insertion.[9, 19–22] Such thiol-disulfide exchange mediated exposure and subsequent insertion of fusion peptide has been reported for other viruses as well.[23–29] On other hand, the presence of disulfide bonds within the membrane active region of reovirus p10 fusion-associated small transmembrane (FAST) proteins[30, 31] and of Ebola virus delta-peptide[32, 33] are quintessential for membrane permeation as demonstrated in recent studies.

Motivated by these studies, we investigate the role of disulfide bond in regulating membrane partitioning of Hepatitis A virus (HAV) 2B protein.[34] Compared to 2B proteins of other picornaviruses, the HAV-2B is unusually longer and shares limited (< 20 %) sequence similarity with them.[35, 36] It plays a vital role in membrane remodelling [37] and viral replication [38], but does not participate in calcium homeostasis or host membrane trafficking [36]. The HAV-2B protein is mainly localized in endoplasmic reticulum membrane and partly in mitochondrial, golgi bodies and plasma membrane.[39] The membrane active part, 60 amino acids long, located at C-terminal region of HAV-2B protein[39], is characterized by presence of multiple cysteine residues. Experimental demonstrations based on biophysical techniques and biochemical assays in membrane mimicking conditions indicated an *α* – helical conformation exhibiting lipid type and composition dependent membrane permeabilizing property.[39] In our previous study[40] based on extensive all-atom MD simulations, we provided insight into HAV-2B peptide induced membrane response as a function of lipid type and composition. The simulations elucidated how the SS-free state of the peptide could sense membrane topography in the form of lipid packing defects and subsequently partitioned into model POPC bilayer, thereby inducing membrane destabilization. We also reported that presence of cholesterol significantly reduced lipid packing defects so as to mitigate peptide recognition and subsequent partitioning into cholesterol-rich membranes. In the current study, we extend our earlier work by considering how chemical changes within the peptide can affect HAV-2B partitioning into model POPC membranes. The presence of multiple cysteine residues in membrane active region of HAV-2B and its localization preference on ER membrane, the site of thiol-disulfide exchange activity, may promote formation of disulfide bonds.

The system of our choice, HAV-2B peptide has three cysteine (C11, C47 and C52) residues. Two out of these three residues may form a SS-bond between them depending on spatial proximity, while the other remains free, resulting in three possible SS-linked states of HAV-2B peptide, denoted as SS11-47, SS11-52 and SS47-52. We perform 500 ns long MD simulations in each of these SS-bonded states and compare with the SS-free HAV-2B peptide to understand how introduction of disulfide bonds may affect partitioning into POPC membrane. The SS-linkage induces shrinking of peptide conformation as well as distortion of its α-helical hairpin geometry in all three states. Further, the presence of disulfide bond restricts insertion and partitioning of HAV-2B peptide into lipid packing defects, subsequently reducing membrane destabilization, unlike that of SS-free state. The disulfide bond thus regulates the membrane active property of the viral peptide. The present findings may have potential implications in drug designing, targeting the HAV-2B protein by promoting disulfide bond formation within its membrane active region. Such therapeutic applications have been implemented or under consideration for treatment of HIV and coronavirus infections.[9, 41]

## 2. Methods

The HAV-2B membrane active viral peptide is 60 amino acids long and harbours three cysteine (C11, C47 and C52) residues (Fig 1). These cysteine residues may remain in reduced thiol state or in oxidized state, where two such residues are bonded through a disulfide linkage, resulting in three possible SS-bonded states of HAV-2B peptide denoted as SS11-47, SS11-52 and SS47-52. Owing to unavailability of experimentally determined structure, the membrane active region of HAV-2B peptide has been modelled in a recent study [39] and further refined through extensive molecular dynamics (MD) simulations in water, details of which are provided in our earlier study[40]. We consider representative MD snapshots of SS-free state of HAV-2B peptide in water to model the SS-bonded states. The snapshots are so chosen such that a pair of cysteine residues in close proximity (~ 5 Å) can form SS-bond between them. The modelled disulfide bonded peptides are further energy minimized over 5000 steps using the conjugate gradient algorithm in NAMD2.10 [42]. We place these energy minimized SS-bonded peptides close (~ 15 Å) to the upper leaflet of previously equilibrated 1-palmitoyl-2-oleoyl-sn-glycero-3-Phosphatidylcholine (POPC) bilayer of surface area ~ 100 × 100 Å^2^, comprising of 147 lipid molecules per leaflet. Sufficient water molecules and counter ions are added to achieve a salt concentration of 0.15 M.

**Fig 1.**
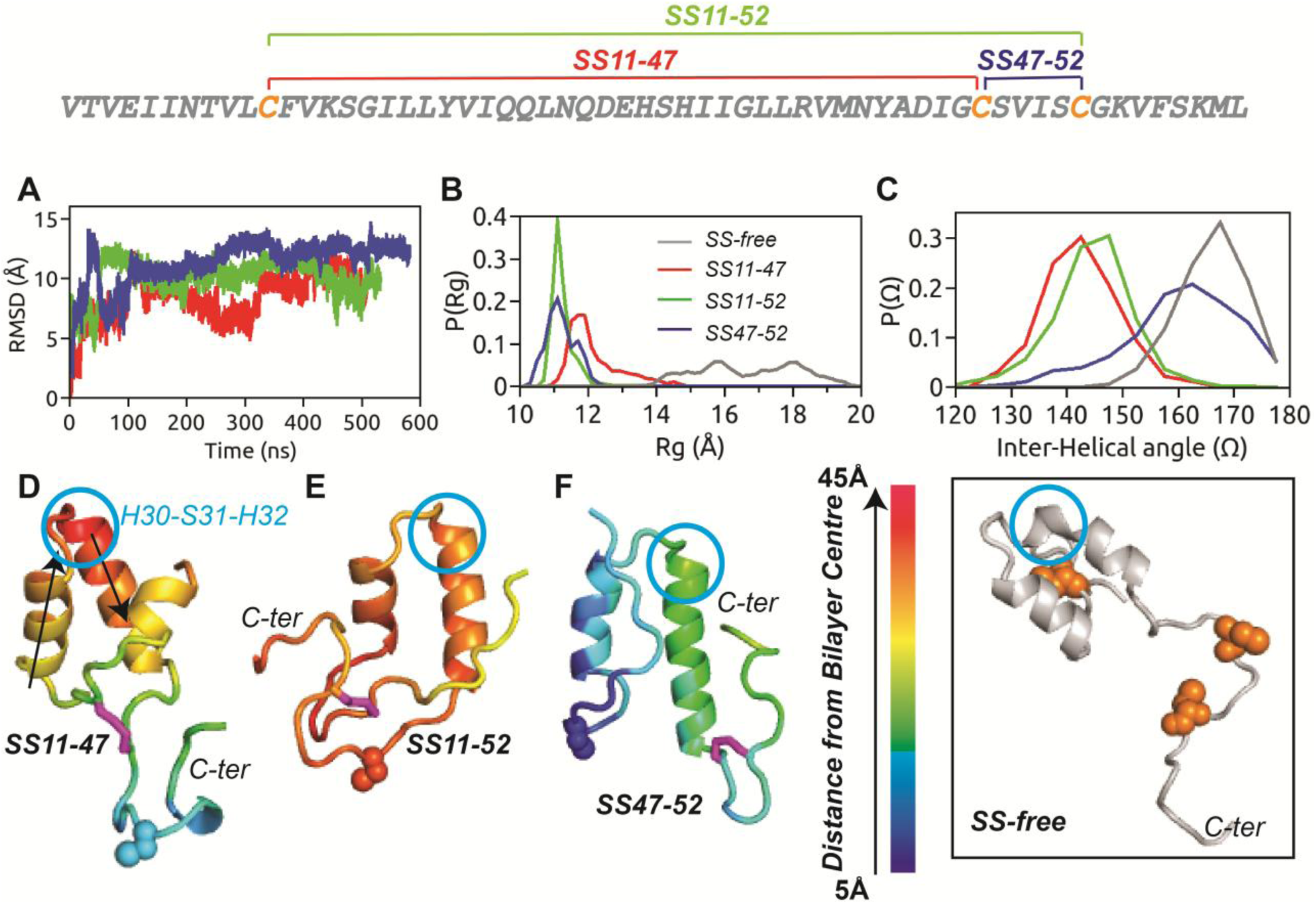
The primary sequence of HAV-2B peptide is shown with the three cysteine residues highlighted in orange, which can form the three possible SS-bonded states. (A) Cα-RMSD of HAV-2B peptide in different SS-bonded states with respect to corresponding initial configurations, showing flat trajectory over the last 200 ns. The equilibrium distributions of (B) radius of gyration, *P*(*R*_*g*_) and (C) inter-helical angle, *P*(Ω) of disulfide bonded states of HAV-2B peptide: SS11-47 (red), SS11-52 (green) and SS47-52 (blue). The *P*(*R*_*g*_) and *P*(Ω) of SS-free HAV-2B peptide (grey) is shown for comparison. The final snapshots of (D) SS11-47, (E) SS11-52 and (F) SS47-52 peptides are illustrated with colour codes representing insertion depth into POPC bilayer; the smaller the value, the deeper is the insertion. The SS-bond is shown in stick representation (magenta), while the cysteine in reduced thiol state is shown in spheres. The arrows in (D) indicate the vectors corresponding to helical axes, used for calculating the inter-helical angle, Ω. The residue triad, H30-S31-H32 which undergoes conformational transition upon inclusion of disulfide bond is indicated by blue circle. The inset shows the largely extended conformation of SS-free HAV-2B peptide and the position of cysteine residues (spheres in orange).

MD simulations for each of these systems (see SI Table S1) are performed in NAMD2.10 [42] using modified TIP3P [43] water model, CHARMM36m [44] and CHARMM36 [45] force field parameters for the peptide and the lipid molecules, respectively. All systems are energy minimized for 10,000 steps. The simulations are carried out using periodic boundary conditions in NPT ensemble at 1 atm pressure and 310K with a time step of 2 femto-seconds. The van der Waals interactions are smoothly truncated beyond 12 Å, by a forced-based switching function between 10 Å and 12 Å, while Particle mesh Ewald fast Fourier transform is used for electrostatic interactions. The peptide heavy atoms are additionally subjected to positional restraints, which are gradually decreased over six cycles of equilibration of 500 ps each to ensure relaxed starting configurations of SS-bonded states. The production runs are performed for over 500 ns and further analysis on equilibrated trajectories are executed using Visual Molecular Dynamics (VMD) [46], MEMBPLUGIN [47], Packmem[48] and in-house Fortran codes.

## 3. Results

The MD simulation of disulfide free HAV-2B peptide in water from our previous study[40], indicate close spatial proximity of cysteine residues, namely C11, C47 and C52, which may favour formation of possible SS-bonded states of the peptide denoted as SS11-47, SS11-52 and SS47-52. The distribution of pairwise SS-distance of disulfide free HAV-2B peptide in water (see SI, Fig S1) is multi-modal with several peaks spread between 3 Å and 20 Å. Relevant MD snapshots with SS-distance < 5 Å are chosen to build respective SS-bonded states, energy minimized and further subjected to restrained MD equilibration protocol as described in the Methods section. We perform 500 ns long all atom MD simulations for each of these states in model POPC membrane. The root mean square deviation (RMSD) plots in Fig 1A based on *C-α* atoms with respect to corresponding initial configurations show structural re-arrangements within the first 300 ns, which we consider as the equilibration period of the SS-bonded peptides in POPC bilayer. The RMSD being flat beyond 300 ns, further analysis is performed on the equilibrated trajectories of last 200 ns. We first present the conformational preferences and the membrane binding mode of different disulfide bonded states. We follow this up with SS-bonded peptide induced membrane response and the role of interfacial packing defects in peptide partitioning. Finally, we elucidate how introduction of the disulfide bond may regulate membrane active property of HAV-2B peptide.

### 3.1 Conformational Preferences

The molecular dimension of HAV-2B peptide is characterized by probability distribution of radius of gyration, *P*(*R*_*g*_) in Fig 1B, computed as the average distance of *C-α* atoms from their centre of mass over the equilibrated trajectories. The SS11-47 and SS47-52 peptides explore conformational states with *R*_*g*_ varying between 11 Å - 14 Å, resulting in broad *P*(*R*_*g*_), in contrast to the sharp distribution of SS11-52 peptide, with ⟨*R*_*g*_⟩~ 11 Å. The presence of SS-bond results in compact conformations compared to the largely extended SS-free state of HAV-2B peptide (see inset Fig 1) in POPC membrane as indicated by *P*(*R*_*g*_) in Fig 1B. The final snapshots illustrating the conformational preferences of the three SS-bonded states of HAV-2B peptide shown in Fig 1D-F, also indicate the location of disulfide bonds. For instance, the SS-linkage connects the N- and the C-terminal tails in SS11-47 (Fig 1D) and SS11-52 (Fig 1E) peptides, while, the linkage is confined within the C-terminal tail of SS47-52 peptide. Despite the differences in location and connectivity, these disulfide bonds act as a constraint reducing the overall conformational fluctuations of the peptide.

We also characterize the conformational preference in terms of the helical arrangement of the hairpin structure through inter-helical angle, Ω. The vector between C-α atoms of L18 and I22 represents the first helical axis, while that of L36 and M40 represents the second, Ω being the angle between them (see Fig 1D). The equilibrium distributions, *P*(Ω) of the SS-bonded states of HAV-2B peptide are shown in Fig 1C. The *P*(Ω) of SS11-47 and SS11-52 peptides are overlapping with ⟨Ω⟩ lying between 140° - 150°, resembling a “*boomerang*” conformation. In contrast, the distribution of inter-helical angle of SS47-52 peptide about ⟨Ω⟩ ~ 160° is quite similar to that of SS-free state (⟨Ω⟩ ~ 170°) resembling a near anti-parallel helical conformation characteristic of a hairpin structure. The SS-linkage between the two terminals thus controls the inter-helical angle in a way so as to distort the hairpin geometry of both SS11-47 and SS47-52 states; whereas the geometry is almost preserved when the linkage is confined within the C-terminal tail of SS47-52 peptide.

The secondary structure of disulfide bonded HAV-2B peptides are calculated using the STRIDE [49] algorithm implemented in VMD [46]. The residue-wise secondary structure percentage (see SI, Fig S2) represents the population of different structural elements explored by each residue during the course of simulation. The residues I17 to L25 and H30 to Y42 form the two *α*-helices of the hairpin motif of the SS-bonded states. The overall secondary structure percentage remains qualitatively similar across different SS-bonded states. The N- and C-terminal tail residues are predominantly characterized by *“turn”* or random *“coil”* conformations, with few residues showing minor populations of *α*-helix, *β*-sheets, and *β* - strands.

We also compare the changes in secondary structure population of the cysteine residues in different SS-bonded states (Fig 2A-C) with that of SS-free state. C11 (Fig 2A) predominantly exhibits random *“coil”* conformation in SS-free as well as in SS47-52 states. But whenever, C11 is involved in SS-bond formation as in SS11-47 and SS11-52 states, the propensity to adopt random *“coil”* conformation decreases with an increase in *“turn”* like secondary structural element. Similarly, random *“coil”* conformation of C47 (Fig 2B) is prevalent in SS-free and SS11-52 states, but enhanced population of *“turn”* is observed in SS11-47 and SS47-52 states.

**Fig 2.**
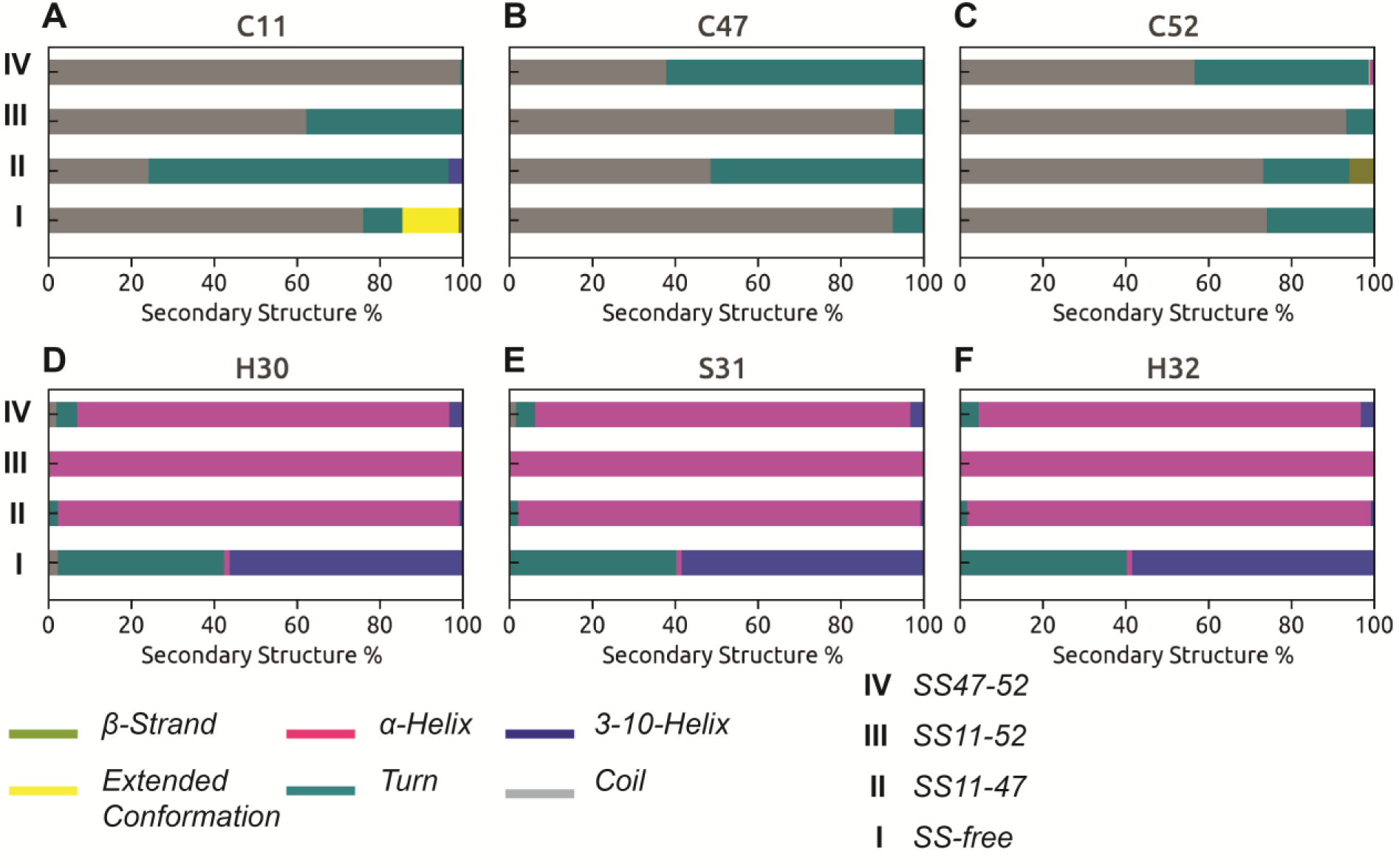
Percentage population of different secondary structural elements accessible to (A) C11, (B) C47, (C) C52, (D) H30, (E) S31 and (F) H32 in SS-bonded and -free states of HAV-2B peptide.

Our observations indicate that the cysteine residues upon being involved in SS-bonding generally show a shift in conformational equilibrium towards *“turn”* like secondary structure component from random *“coil”* conformation, the only exception being increased *“coil”* propensity for C52 (Fig 2C) in SS11-52 state. The presence of disulfide bond significantly alters the secondary structure propensity of a triad of residues, namely, H30-S31-H32 (see Fig 2D-F), the location of this triad is indicated in Fig 1D-F. The residue triad forms part of the inter-helical flexible linker in SS-free state and shows transition between *“turn”* and 3_10_-helices. In contrast, these residues adopt an *α*-helical conformation all throughout the SS-bonded states (Fig 2D-F). The change in conformational equilibrium of H30-S31-H32 from flexible *“turn”* / 3_10_-helical to more rigid *α*-helical conformation possibly accounts for the observed deviation in inter-helical angles (Fig 1C) of SS-bonded states from that of SS-free state.

### 3.2 Peptide-Bilayer Mode of Binding

We observe significant differences in mode of peptide binding to POPC bilayer depending on disulfide connectivity (Fig 3). The SS11-47 peptide interacts with POPC membrane (Fig 3A) through its C-terminal region. The α-helical hairpin motif orients itself almost parallel to membrane normal (z-direction) and remains solvent exposed including the N-terminal tail. The localization of peptide residues on bilayer is quantified through its insertion depth, calculated as the distance of centre of mass of each peptide residue from bilayer centre along the z-direction. The final snapshots of the SS-bonded peptides in Fig 1D-F are colour coded according to residue insertion depths: The smaller the value, the deeper is the insertion. Fig 1D indicates that the C-terminal residue stretch V49 – F56 of SS11-47 peptide is located in close proximity (~ 15 Å) to bilayer centre indicating insertion, in contrast to the α-helical hairpin motif and the N-terminal tail, which remain far (~ 40 Å) from membrane interior. The SS11-52 peptide hovers close to membrane surface but fails to form any stable contacts with POPC membrane (Fig 3B) within our simulation time-scale of 500 ns. As a consequence, the peptide residues remain far (~ 40 Å) from bilayer centre (Fig 1E). The binding mode of SS47-52 peptide with POPC membrane is shown in Fig 3C. Unlike SS11-47 peptide, the α-helical hairpin motif orients itself parallel to bilayer resulting in enhanced contact surface area. The SS47-52 peptide interacts with the membrane by complete insertion of N-terminal helix (I17 – L25) and the preceding tail, the average insertion depth being ~ 10 – 15 Å (Fig 1F). The C-terminal helix (H30 – Y42) of the hairpin motif and the succeeding tail region, although solvent exposed, are located close to the POPC headgroups.

**Fig 3.**
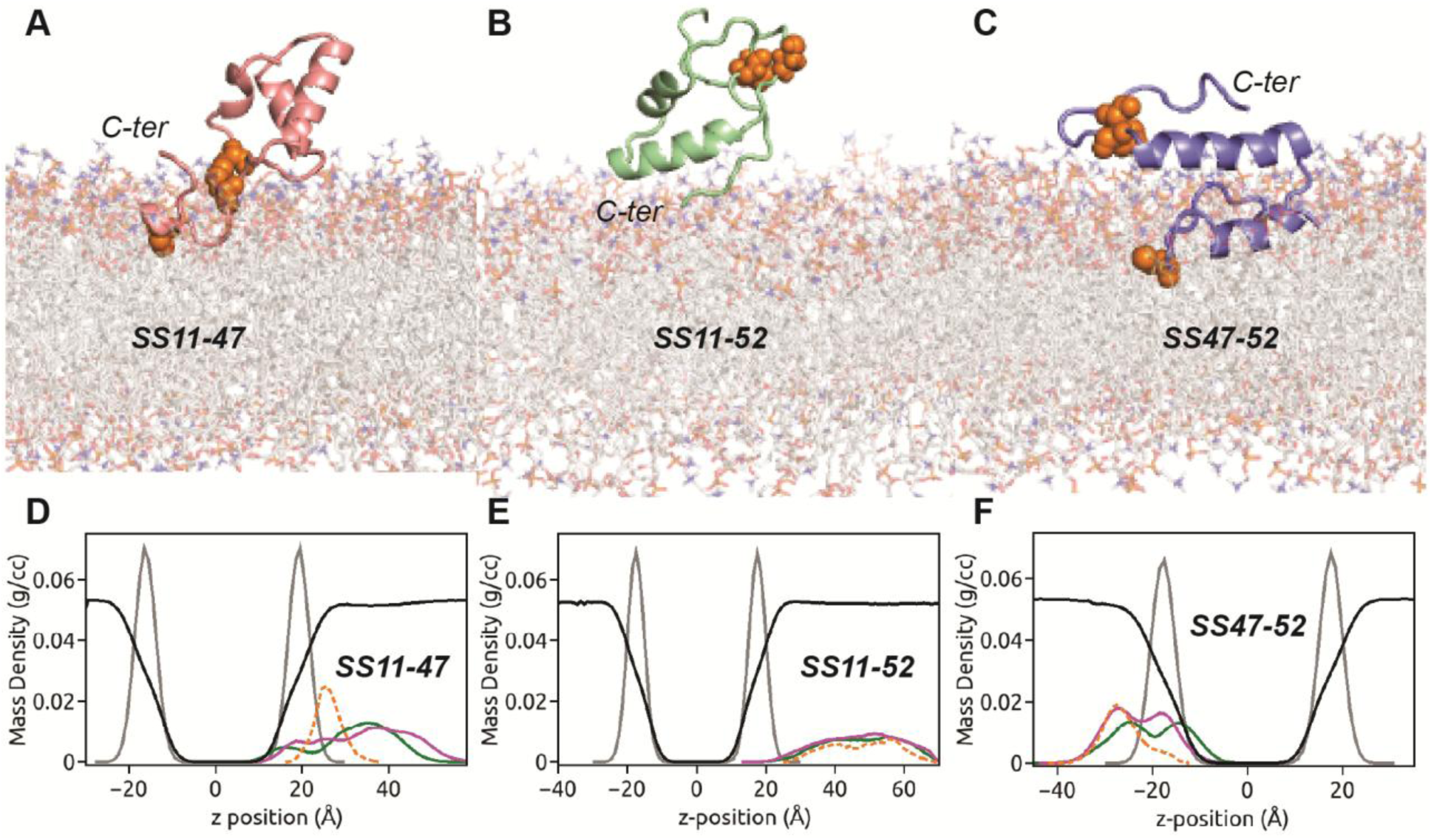
The binding modes of (A) SS11-47, (B) SS11-52 and (C) SS47-52 peptides with POPC bilayer are shown. The cysteine (both SS-linked and free) residues are represented as orange spheres, while hydrating water is not shown for clarity. The mass density profiles along the bilayer normal (z-direction) of (D) SS11-47, (E) SS11-52 and (F) SS47-52 peptides in POPC bilayer. The density profiles comprising of hydrophobic (green) and hydrophilic (magenta), SS-bonded cysteine (orange) residues of peptide, phosphate headgroups (grey) and water (reduced by factor of 10, black) are indicated.

The peptide-membrane mode of binding influences the degree of membrane induced partitioning of peptide. The partitioning of SS11-47 peptide in POPC bilayer is illustrated through density profile of peptide atoms along the z-direction as a function of distance from membrane centre (Fig 3D), calculated over the equilibrated trajectory. As bulk of the SS11-47 peptide remains in water, the density profiles of constituent hydrophobic and hydrophilic residues are overlapping with peak positions in solvent. The peptide density profiles also show little overlap with that of lipid headgroups owing to insertion of few C-terminal residues. Since the SS11-52 peptide fails to insert into membrane, the corresponding density profile in Fig 3E is entirely positioned in solvent rarely overlapping with lipid headgroups. Although, we initially place the SS47-52 peptide close to the top leaflet, during the course of simulation, the peptide leaves the central simulation box from the top to re-enter from the bottom and localize on the lower leaflet. This results in the density profile of SS47-52 peptide to overlap with that of lipid headgroups from the lower leaflet, as indicated in Fig 3F. The SS47-52 peptide partitions deep into the membrane milieu with segregation of hydrophobic and hydrophilic density peaks towards bilayer centre and headgroups, respectively, acquiring partly facially amphiphilic conformation in POPC bilayer. Both hydrophobic and hydrophilic density profiles being bimodal, a second peak is observed in solvent close to the headgroup-water interface. This bimodal nature of density profiles is attributed to the observed horizontal binding mode of SS47-52 (Fig 3C), characterized by membrane embedded N-terminal helix and surface adsorbed C-terminal helix near the POPC headgroups.

The horizontal orientation of SS47-52 peptide parallel to membrane surface is quite similar to the observed binding mode of SS-free HAV-2B peptide, except that in the latter both helices of the hairpin motif and the long stretched C-terminal tail remain more deeply (~5-10 Å from bilayer centre) embedded within the POPC membrane.[40] Thus more number of hydrophobic and hydrophilic residues forms extensive contacts with lipid molecules resulting in enhanced density profile peak intensities of SS-free state (see SI Fig S3) compared to SS47-52 peptide. Further, the segregation of hydrophobic and hydrophilic peak intensities facilitate the SS-free HAV-2B peptide to acquire a strong facially amphiphilic conformation in POPC membrane. Our results strongly demonstrate that disulfide connectivity can regulate the extent of HAV-2B peptide partitioning. The disulfide linkages connecting the N- and C-terminal tails of HAV-2B peptide, as in SS11-47 and SS11-52 states, mitigates peptide partitioning. Whereas, the SS-linkage confined within the C-terminal tail involving the C47-C52 pair, results in peptide partitioning, albeit weaker than the SS-free state.

In this regard, it is also interesting to study the partitioning of cysteine residues. The density peaks of disulfide bonded cysteine pairs in SS11-47 (Fig 3D) and SS47-52 (Fig 3F) states are located at the solvent proximal interface very close to the lipid headgroups. While the other cysteine in reduced thiol state: C52 of SS11-47 (Fig 3A) and C11 of SS47-52 (Fig 3C) peptides remain embedded in the hydrophobic membrane core. This is in agreement to a recent study, which concluded that cysteine residues in reduced thiol state favourably partitions into the hydrophobic membrane milieu rather than at the polar lipid-water interface.[50]

### 3.3 Influences on Membrane Properties

We investigate how membrane properties are affected upon partitioning of SS47-52 peptide in model POPC membrane. To this end, we quantify the flexibility of lipid acyl chains through lipid tail order parameter, 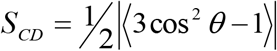 being the angle between C-H bond vector and bilayer normal, computed over the equilibrated trajectories for both saturated (*sn* – 1) and unsaturated (*sn* – 2) acyl chain carbon atoms and shown in Fig 4. Higher the order parameter, the lower is the flexibility of lipid acyl chains and vice-versa. The order parameter values, *S*_*CD*_ of *sn* – 2 (Fig 4A) and *sn* – 1 (Fig 4B) chains of POPC under the influence of SS47-52 peptide partitioning are intermediate between that of control POPC bilayer (without any peptide) and in presence of SS-free HAV-2B peptide. The presence of C47-C52 disulfide linkage enhances the acyl chain flexibility compared to control POPC bilayer, however, the disordering effects are not as strong as in the presence of the SS-free peptide. The enhanced flexibility of lipid tails causes lateral expansion of bilayer, leading to increased surface area-per-lipid (SA/lipid) and reduced bilayer thickness. The effect of SS47-52 peptide on SA/lipid (~ 74 Å^2^) and bilayer thickness (~ 35.6 Å) is also observed to be intermediate between that of SS-free peptide and control POPC bilayers.[40] We illustrate the 2d-thickness profiles (Fig 4C-D) along the membrane xy-plane corresponding to final snapshots (Fig 4E-F) of SS47-52 and SS-free peptide-membrane systems. The lateral thickness profile reflects membrane thinning localized around the SS47-52 peptide, with thickness varying between ~ 30 Å at insertion site to ~ 40 Å elsewhere. In contrast, the SS-free state of HAV-2B peptide induces uniform global thinning of POPC bilayer, the effect being more pronounced (~ 30-35 Å) compared to the SS47-52 peptide. Owing to the reduced partitioning of HAV-2B peptide in presence of SS-bond, the membrane perturbation effects are also significantly reduced.

**Fig 4.**
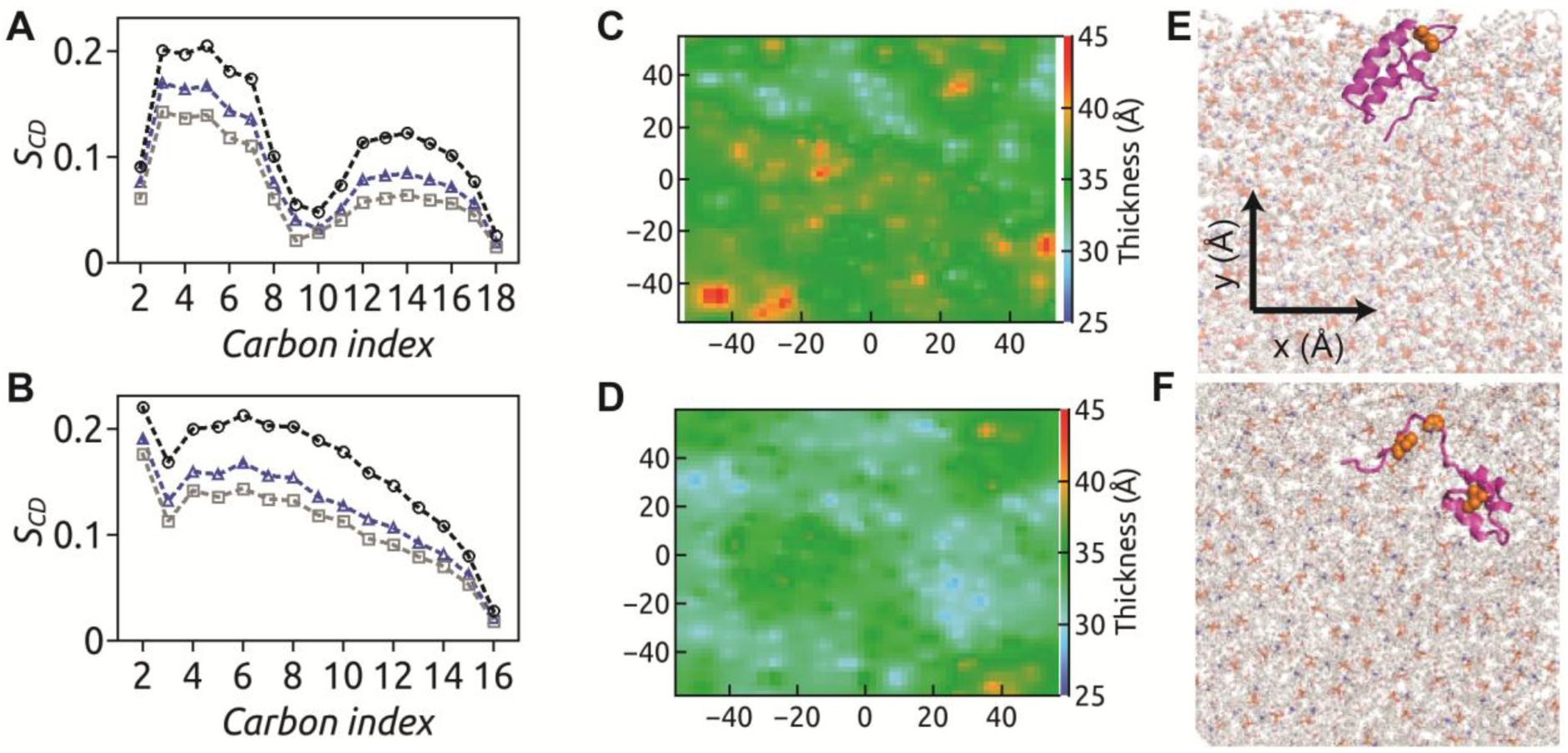
The order parameter, *S*_*CD*_ of (A) *sn* – 2 and (B) *sn* –1acyl chains of POPC corresponding to SS47-52 (blue triangle), SS-free peptide (grey square) and control POPC (black circle) systems are shown. The membrane thickness maps upon interaction of (C) SS47-52 and (D) SS-free states of HAV-2B peptide with POPC bilayer are generated considering inter-leaflet P-P distance with 2Å resolution along the xy-plane. The localization of (E) SS47-52 and (F) SS-free peptide on POPC bilayer are shown. The membrane thinning effect is localized around the insertion site of SS47-52 peptide, while more pronounced uniform global thinning effect is observed upon SS-free HAV-2B peptide partitioning.

### 3.4 Distribution of Lipid Packing Defects

The membrane response is not only restricted to lipid acyl tails, but also extend to the bilayer-water interface in the form of lipid headgroups packing. This interfacial region is characterized by transient exposure of membrane hydrophobic core to hydration layer, leading to lipid packing defects which act as binding hotspots for peptides.[51–60] These lipid packing defects are qualitatively characterized into “Deep” or “Shallow” depending on the relative depth of the defect site with respect to the nearest glycerol backbone and further quantified by area (*A*) of the defect site, following the standard protocol using Packmem [48]. The distribution of defect sites based on their size exhibit single exponential decay, *P*(*A*) = *B* exp(– *A* П). The decay constant, П, characteristically describes the membrane topology in terms of interfacial defects: The higher the value of П, the more abundant are large defect sites.

The size distribution of “Deep” defect sites is shown in SI, Fig S4A, truncated to include upto moderate (~ 100 Å^2^) sized defects with occurrence probability > 10^−4^. The relative population of different defect sizes in presence of SS-bonded peptides are almost similar following a single-exponential behaviour within the range considered but deviates beyond defect sizes > 60 Å^2^. The calculated defect area constant, П_*Deep*_, from single exponential fits, summarized in Table S2, indicate similar values varying between 10 – 12 Å^2^. The size distributions of “Shallow” defects, shown in SI, Fig S4B, are also similar for all SS-bonded peptides with distributions, П_*Shallow*_ varying between 15 – 17 Å^2^ (see Table S2). Upon comparing the size *P*(*A*) of both “Deep” and “Shallow” defects, we observe that the number of defect sites as well as the occurrence of larger defect sites increase in presence of SS-bonded peptides with respect to control POPC bilayer. However, this abundance of defect sites is much more pronounced under the influence of SS-free HAV-2B peptide, as also evident from higher defect area constants listed in Table S2.

The defect size distributions *P*(*A*) reveal the abundance of varying defect sizes that accumulate over time, but do not quantify the extent of defects in a given frame. So we calculate the total defect area in a given frame by adding the individual areas of all “Deep” (or “Shallow”) defect sites in the frame and normalize by the area of leaflet, to define the “Deep” (or “Shallow”) defect area fraction *f*_*Deep*_ (or *f*_*Shallow*_) in the given frame. It provides a measure of how much leaflet area is covered by “Deep” (or “Shallow”) lipid packing defects in a given frame. The distribution of “Deep” defect area fraction, *P*(*f*_*Deep*_) due to presence of SS-bonded peptides in POPC bilayer is shown in Fig 5A. The *P*(*f*_*Deep*_) of SS11-47 and SS11-52 are single-peaked around defect area fractional values ~ 0.02. The overlapping distributions indicate that the extent of “Deep” defects is similar for both these “*non-partitioning*” SS-bonded peptides but more compared to the control system. On contrary, *P*(*f*_*Deep*_) of SS47-52 and SS-free state are broad and have significant overlap at higher defect area fraction values, *f*_*Deep*_ ~ 0.04. This implies that large amount of leaflet area is covered by “Deep” defects in presence of these peptides. Since both the helices of SS-free peptide partition into POPC bilayer, the defect area fraction is slightly higher compared to SS47-52, where only the N-terminal helix partitions. The extent of “Shallow” defects, similar in all SS-bonded systems due to overlapping *P*(*f*_*Shallow*_) (Fig 5B) with peak around 0.04, are intermediate between that of control POPC and SS-free system. Unlike the “Deep” defects, the “Shallow” defects remain unaffected by partitioning of SS47-52 peptide. While “Deep” defects are affected by partitioning of bulky hydrophobic residues, “Shallow” defects are known to be influenced by presence of short chain hydrophobic amino acids, as reported for *α*–Synuclein.[54, 56] The N-terminal helix of hairpin motif and the preceding tail harbours few such small hydrophobic residues, while the majority of them reside at the C-terminal helix and the succeeding tail. Owing to this, the partitioning of N-terminal helix of SS47-52 does not affect the “Shallow” defects. On contrary, both the helices and the C-terminal tail of SS-free peptide being involved in membrane partitioning, enhances the “Shallow” defects.

**Fig 5.**
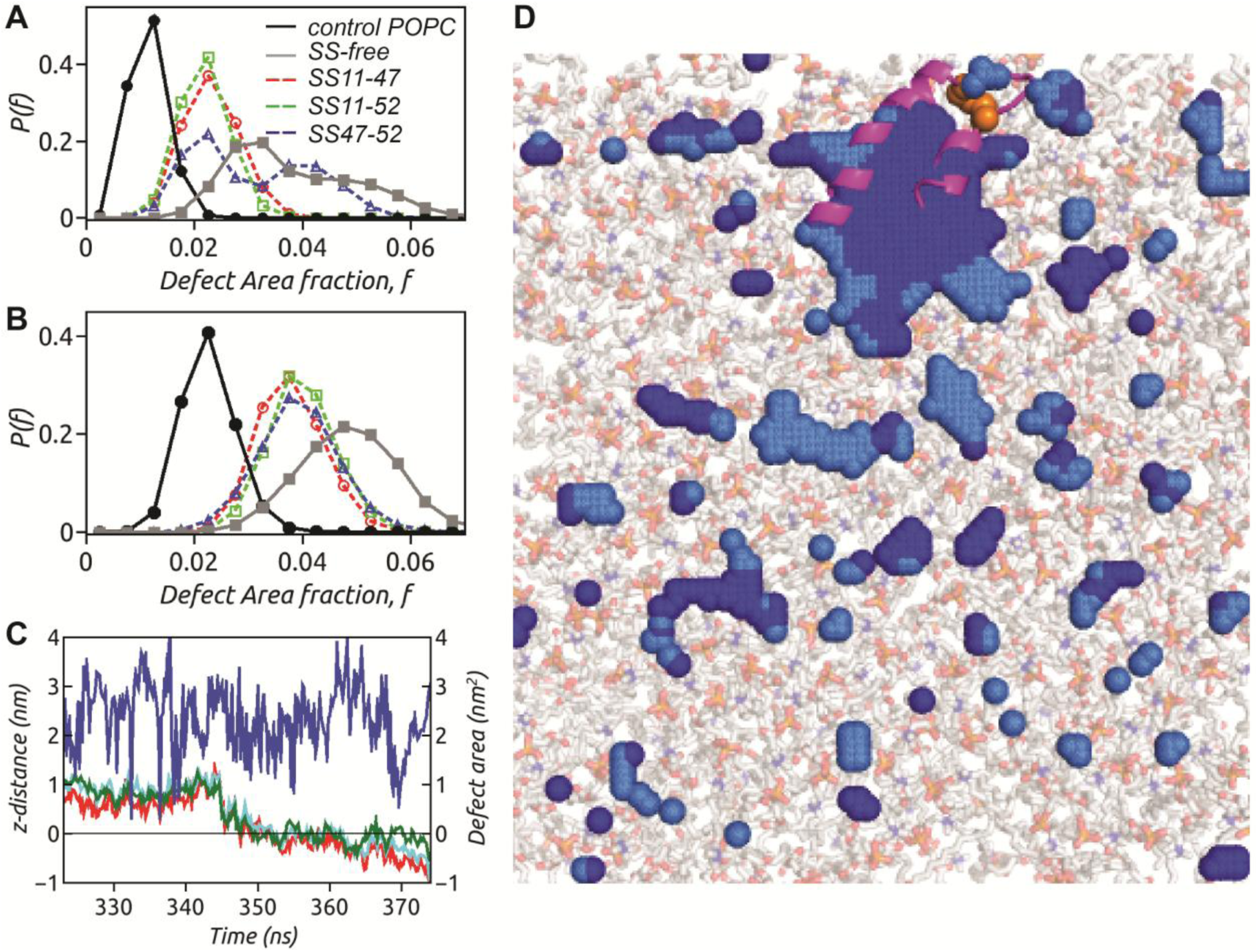
The distribution of defect area fraction *P*(*f*) corresponding to (A) “Deep” and (B) “Shallow” provide insight into the extent of defects in a given frame relative to leaflet area. The open symbols with dotted line represent SS11-47 (red circle), SS11-52 (green square) and SS47-52 (blue triangle) systems, while solid symbols with solid lines indicate SS-free peptide (grey square) and control (black circle) POPC system. (C) The insertion dynamics of C11 (red), L18 (cyan) and L25 (green) of SS47-52 peptide into a co-localized “Deep” defect area (blue). (D) The final snapshot of SS47-52 peptide with its N-terminal helix completely embedded into the large co-localized “Deep” defect (dark blue) surrounded by “Shallow” defects (light blue). The SS-linked cysteine residues (orange spheres) fail to insert into the co-localized defect.

### 3.5 Sensing of Lipid Packing Defects

Our previous study indicated that the SS-free HAV-2B peptide senses membrane topography in the form of lipid packing defects, inserts into such defects and subsequently partitions into POPC bilayer.[40] In this section, we elucidate the mechanism of partitioning of SS47-52 state of the viral peptide. We consider a representative set of residues, C11 (red), L18 (cyan) and L25 (green) of SS47-52 peptide (Fig 5C), which undergo deep insertion into POPC membrane. The insertion dynamics of these residues are monitored from the individual distance (z-distance) of residue centre of mass from the average level of C2 atoms of glycerol moieties in POPC molecule, along the z-direction. A negative value of z-distance implies insertion below the average C2 level. Simultaneously, we track the appearance of any underlying “Deep” lipid packing defect that is co-localized with these residues. In the process, we identify a single large co-localized “Deep” defect in vicinity of these residues. The defect area fluctuating around 250 Å^2^ drives the residue insertions at around 350 ns, following which the co-localized “Deep” defect area momentarily increases to 400 Å^2^ to accommodate the bulky hydrophobic side-chains. The insertion of SS47-52 peptide into the co-localized “Deep” defect is illustrated through a representative snapshot in Fig 5D. The horizontal orientation of the inserted N-terminal helix of the hairpin motif stabilizes the large “Deep” defect ~ 270 Å^2^. This single large defect contributes to the SS47-52 peptide induced enhanced “Deep” defect area fraction *P*(*f*_*Deep*_) in Fig 5A. These discrete residue insertion events following the appearance of co-localized defects suggest sensing of lipid packing defects similar to the SS-free peptide

## 4. Discussion

The present study provides insight into the effect of disulfide bond on HAV-2B peptide structure and partitioning into membrane. The inclusion of SS-bond not only results in a compact conformation but also changes the inter-helical angle, Ω, resulting in deviation from hairpin conformation in comparison to SS-free state. The anti-parallel hairpin arrangement of α-helices is known to be essential for membrane partitioning of viral peptide as reported for Influenza virus hemagglutinin[61] and Ebola virus delta peptide[33]. The change in inter-helical angle, which in turn modulates helix packing interactions along with the compact conformation of SS-bonded states control the accessible surface area (ASA) of the peptide (Fig 6A). The hydrophobic residues being crucial in regulating HAV-2B peptide partitioning, we calculate the contribution of the same to ASA in presence and absence of the SS-bond. The hydrophobic accessible surface area (*A*_*H*_) is significantly high (~ 3000 Å^2^) in SS-free state, followed by SS47-52 state (2300 Å^2^). The disulfide bond interconnecting the N- and C-terminal tails further reduces the hydrophobic exposure with *A*_*H*_ < 2000 Å^2^ observed in SS11-47 and SS11-52 peptides, which thus fail to partition into POPC bilayer. The SS-free state of HAV-2B peptide with maximum hydrophobic exposure partitions deep into membrane milieu, while the partitioning ability is partly compromised with reduced hydrophobic exposure upon inclusion of SS-bond between C-terminal cysteines, C47-C52.

**Fig 6.**
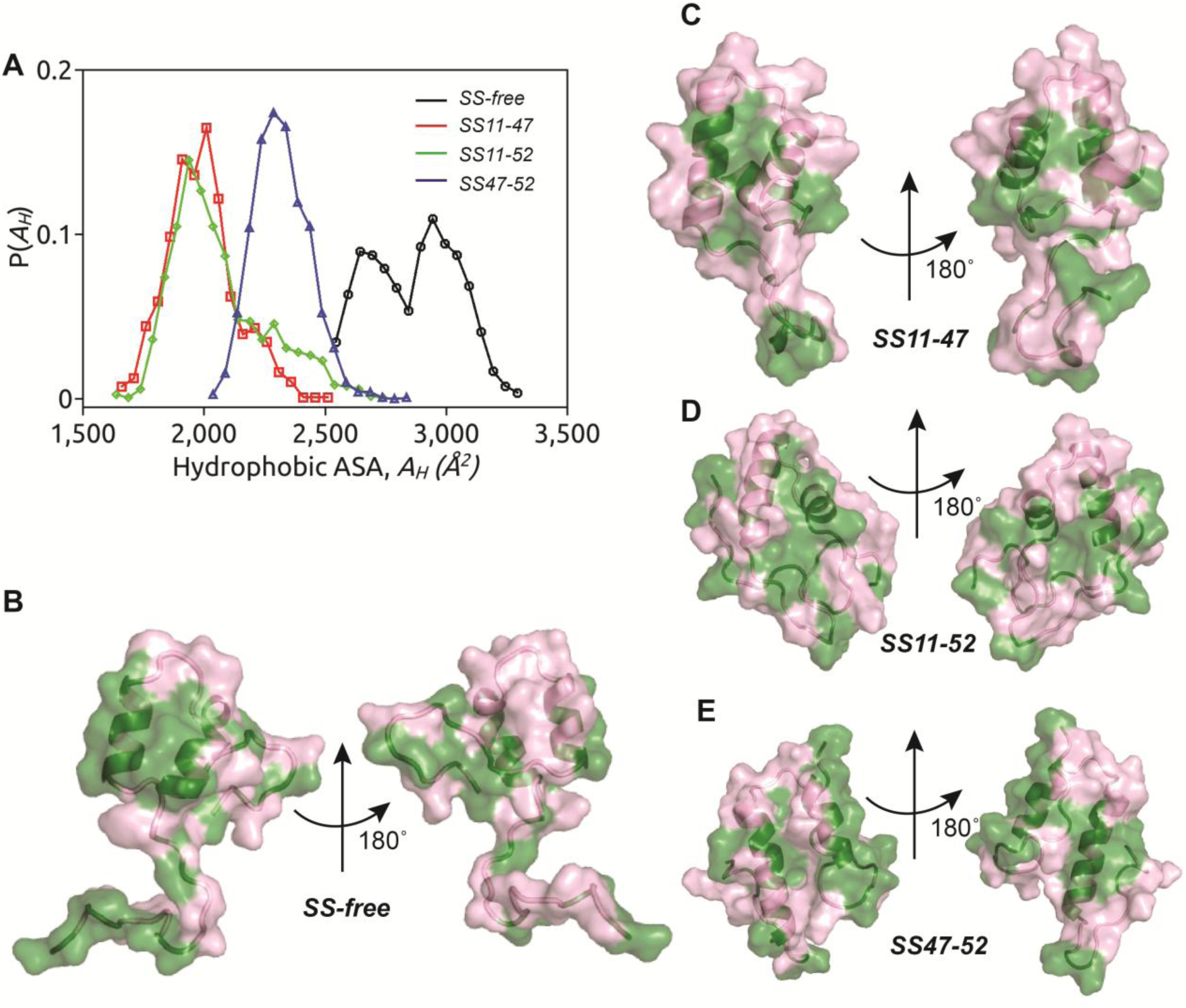
(A) The distribution of hydrophobic accessible surface area (ASA), *P*(*A*_*H*_) is shown for SS-free (black), SS11-47 (red), SS11-52 (green) and SS47-52 (blue) states of HAV-2B peptide. The solvent exposed hydrophobic (green) and hydrophilic (magenta) surface area is illustrated on the 3-D structure of HAV-2B peptide in (B) SS-free, (C) SS11-47, (D) SS11-52 and (E) SS47-52 states.

We illustrate the hydrophobic (green) and hydrophilic (magenta) ASA of HAV-2B peptide in SS-free (Fig 6B) and bound states (Fig 6C-E). In SS-free state, the peptide presents an exposed hydrophobic dominated face and a hydrophilic dominated opposite surface (Fig 6B). The conformation of SS-free HAV-2B peptide is such that it acquires a strong facially amphipihilic character upon segregation of hydrophobic and hydrophilic residues. Different membrane active agents including antimicrobial peptides [55, 62, 63], polymers [58, 64–69] and other membrane active molecules [70–72] are known to acquire such amphipihilic conformations upon partitioning into cellular membranes. However, in SS11-47 (Fig 6C) and SS11-52 (Fig 6D) states, the hydrophobic accessible surface area is limited resulting in mitigation of peptide partitioning. On other hand, the segregation of hydrophobic and hydrophilic surfaces is not as discrete as that in SS-free state, leading to partial partitioning of SS47-52 peptide (Fig 6E). The disulfide bond induced conformational changes thus control the exposure of hydrophobic residues, which in turn regulates HAV-2B peptide partitioning. As a consequence the SS-bonded peptide induced membrane responses, in terms of lipid tail disordering, membrane thinning and abundance of interfacial packing defects, are mild compared to the SS-free state. In a way, the disulfide bond regulates the membrane active property of HAV-2B peptide, such that the membrane destabilizing effects of this viral peptide are significantly reduced.

Evidence of thiol-disulfide redox status dependent exposure of hydrophobic patches facilitating peptide partitioning is well documented through experimental investigations on different membrane active agents like antimicrobial peptides[10, 11, 13, 17] and viral fusion peptides[21, 23–29, 31]. This importance of disulfide bond in mediating viral peptide partitioning and subsequent entry into host cells is currently being explored to design antiviral agents. For instance, reduction of disulfide bond by PDI being pre-requisite for HIV entry, designing inhibitors targeting this process interferes with the virus / cell fusion mechanism.[9, 19–21] Similar efforts are in progress to develop therapeutics against coronavirus infection.[41] The present findings indicate that promoting disulfide bond formation within the membrane active HAV-2B peptide may have potential implications in designing antiviral agents to combat HAV infection.

## 5. Conclusion

The presence of multiple cysteine residues in the membrane active region of HAV-2B peptide indicates the possibility of three SS-bonded states of the peptide. In the present work, we elucidate the role of disulfide bond in HAV-2B peptide partitioning. The SS-linkage induces shrinking of peptide conformation as well as distortion of its α-helical hairpin geometry, resulting in reduced hydrophobic exposure. Depending on disulfide connectivity, the partitioning of HAV-2B peptide is completely or partly abolished and subsequently reduced membrane remodelling effects are observed in comparison to SS-free state. The disulfide bond thus regulates the membrane active property of the viral peptide. These results may find potential applications in drug designing approaches against HAV infection.

## Author Contributions

SS, MB and SV designed the project. SS performed the simulations and carried out the analysis. All authors contributed to writing and reviewing the manuscript.

## Acknowledgement

All simulations in this work have been carried out on supercomputing facility Nandadevi cluster at The Institute of Mathematical Sciences, Chennai, India.

